# A conserved D/E-P motif in the nucleotide binding domain of plant ABCB/PGP-type ABC transporters defines their auxin transport capacity

**DOI:** 10.1101/2020.05.08.084087

**Authors:** Pengchao Hao, Jian Xia, Jie Liu, Martin diDonato, Konrad Pakula, Aurélien Bailly, Michal Jasinski, Markus Geisler

**Affiliations:** University of Fribourg, Department of Biology, CH-1700 Fribourg, Switzerland; Department of Plant Molecular Physiology, Institute of Bioorganic Chemistry, Polish Academy of Sciences, Noskowskiego 12/14, PL-61-704 Poznan, Poland; Institute for Plant and Microbial Biology, Zollikerstrasse 107, 8008 Zurich, Switzerland; Shanghai Jiaotong University, School of Agriculture and Biology, Shanghai, China; NanoBioMedical Centre, Adam Mickiewicz University, Umultowska 85, PL-61-614 Poznan, Poland; Department of Biochemistry and Biotechnology, Poznan University of Life Sciences, Dojazd 11, 60-632 Poznan, Poland

**Author notes:** Corresponding author: Markus Geisler.

**Keywords:** ABC transporter, ABCB, PPIase, FKBP, Twisted Dwarf1, auxin transport

## Abstract

Auxin transport activity of ABCB1 was suggested to be regulated by physical interaction with the FKBP42/Twisted Dwarf1 (TWD1), a *bona fide* peptidylprolyl *cis-trans* isomerase (PPIase), but all attempts to demonstrate such a PPIase activity on TWD1 have failed so far.

By using a structure-based approach we have identified a series of surface-exposed proline residues in the C-terminal nucleotide binding fold and linker of Arabidopsis ABCB1 that do not alter ABCB1 protein stability or location but its catalytic transport activity. P1.008 was uncovered as part of a conserved signature D/E-P motif that seems to be specific for <u>A</u>uxin-<u>T</u>ransporting <u>A</u>BCBs, we now refer to as ATAs. Beside the proline, also mutation of the acidic moiety prior to the proline abolishes auxin transport activity by ABCB1. So far, all higher plant ABCBs for that auxin transport was safely proven carry this conserved motif underlining its diagnostic potential. Introduction of this D/E-P motif into malate importer, ABCB14, increases both its malate and its background auxin transport activity, suggesting that this motif has an impact on transport capacity. The D/E-P1.008 motif is also important for ABCB1-TWD1 interaction and activation of ABCB1-mediated auxin transport by TWD1, supporting a scenario in that TWD1 acts as an activator of ABCB1 transport activity by means of its PPIase.

In summary, our data imply a dual function for TWD1 acting as an ABCB co-chaperone required for ABCB biogenesis and as a putative activator of ABCB-mediated auxin transport by *cis-trans* isomerization of peptidyl-prolyl bonds.

## Introduction

The polar distribution of the plant hormone, auxin, is a unique, plant-specific process and the primary cause for the establishment of local auxin maxima and minima that trigger a plethora of physiological and developmental programs (1-3). The generation, control and flexibility of these gradients require the coordinated action of members of at least three major auxin transporter families: the PIN-FORMED (PIN), the AUXIN1-RESISTANT1 (AUX1)/LIKE AUX1 (AUX1/LAX) and the ABC transporters of the B family (ABCBs) (4-6). In Arabidopsis, so-called long PINs (PIN1, 2, 3, 4 and 7) function as plasma membrane (PM) permeases (5), while cytoplasmic entry over the plasma membrane was shown to be dependent on AUX1/LAX proteins thought to function as high-affinity auxin-proton symporters (7,8). In contrast, a subgroup of <u>a</u>uxin-transporting <u>A</u>BCBs (referred to as ATAs in the following) functions as primary active (ATP-dependent) auxin pumps that are able to transport against steep auxin gradients (9-11).

Out of the 22 full-size ABCB isoforms in Arabidopsis, ABCB1, 4, 6, 14, 15, 19, 20 and 21, were associated with polar auxin transport (PAT) (12-17), however, only for ABCB1, 4, 6, 19, 20, 21, transport activities were confirmed (11-13,15,16,18-20). Interestingly, whereas Arabidopsis ABCB1, 6, 19, 20 and tomato ABCB4 (21) were shown to function as specific auxin exporters, Arabidopsis ABCB4 and 21 (11,13,16,19) and rice (*Oryza sativa)* ABCB14 (22) were suggested to function as facultative IAA importers/exporters. In contrast, Arabidopsis ABCB14 is considered as malate importer (23). Currently, it is not known how many of the remaining 15 full-size ABCBs and 7 half-size ABCBs in Arabidopsis function as well as auxin transporters (see Suppl. Fig. 1).

All characterized auxin transporters have been shown to be regulated on the transcriptional and non-transcriptional level (9,24,25). PM PINs and ABCBs were shown to be regulated by a partially overlapping subset of members of the AGC kinase family (26,27). While protein phosphorylation of the central PIN loop seems to alter both their polarity and transport activity (28), phosphorylation of the so-called ABCB linker has been described to result only in activity changes (26-28).

Further, PINs and ABCBs seem to be also regulated by protein-protein interaction with *cis-trans* peptidylprolyl isomerases (PPIases), which are formed by three evolutionary unrelated families: the cyclophilins, the FK506-Binding Proteins (FKBPs) and the parvulins (9). The cyclophilin, DIAGEOTROPICA (DGT/ CypA), seems to decrease the functionality of PINs and their presence on the PM (29) although underlying mechanisms are far from being understood. Further the parvulin, PIN1At, was shown to affect PIN1 polarity by means of its PPIase activity (30). In contrast, biogenesis of the ATAs, ABCB1, 4,19, was found to depend on the FKBP42, TWISTED DWARF1 ((TWD1)/ ULTRACURVATA 2 (UCU2)). The latter concept is based on the finding that ABCB1,4,19 are retained on the ER and degraded in *twd1* (31,32). This model is further supported by that fact that *twd1* and *abcb1,19* plants reveal widely overlapping but not entirely identical auxin-related phenotypes, including dwarfism and a non-handed helical rotation of the epidermal cell files, designated as epidermal “twisting” (31,32).

TWD1 was shown to interact physically via its PPIase/ FK506-binding (FKB) domain (FKBD) with the C-terminal nucleotide binding fold (NBD2; see Fig. 1) of ABCB1, however, the exact mechanism through which TWD1 regulates ABCB biogenesis and/or auxin transport is not yet known. High-molecular weight FKBPs usually possess PPIase activity that is associated with the FKBD (33,34) and a chaperon activity that is thought to be linked with the tetratricopeptide repeat (TPR) domain (33,34). The latter was demonstrated for TWD1 (35), however, all attempts to demonstrate a PPIase activity on recombinant TWD1 protein have failed so far (36).

**Figure 1:**
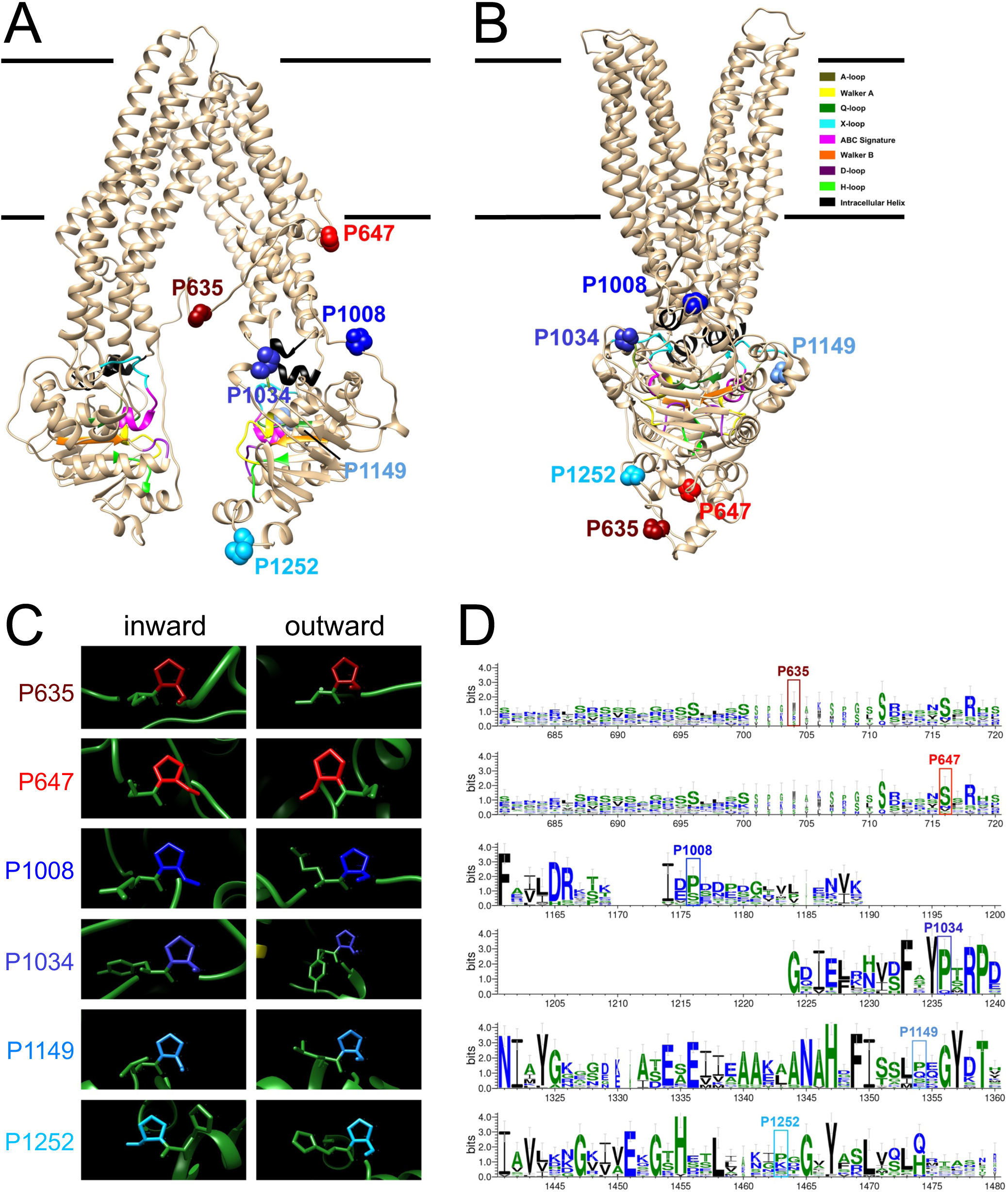
Mapping and conservation of surface-exposed proline residues on Arabidopsis ABCB1. **(A-B)** Surface-exposed prolines of the C-terminal nucleotide binding fold (NBD2) and the linker are indicated as blue and red calottes, respectively, in the in-inward open (**A**) and outward-open conformations (**B)** of Arabidopsis ABCB1 modeled on human PGP/ABCB1 as described elsewhere (49). Color code of functional NBD sequences can be deduced from the legend. **(C)**Close-up analyses of linker and NBD2 prolines indicate that all peptidyl-prolyl bonds are in *trans*. **(D)** Logo representation (WebLogo 3) of linker and NBD2 proline conservation in Arabidopsis after multiple sequence alignment with MUSCLE.

Interestingly, the proposed chaperone-like function for TWD1 during ER to plasma membrane trafficking of ABCBs is in analogy with the functional ortholog of TWD1/FBP42 in humans, FKBP38/FKBP8, that shares with TWD1/FKBP42 a unique mode of membrane-anchoring (37,38). FKBP38 promotes ER to plasma membrane delivery of the cystic fibrosis transmembrane conductance regulator, CFTR/ABCC7 (38). CFTR, belonging to ABCC/ MRP family of ABC transporters, is a chloride channel whose malfunction is responsible for the genetic disorder mucoviscidosis (37). About 90% of all cystic fibrosis cases are associated with a specific single residue deletion, CFTR^ΔF508^ (38,39). This mutation results in a folding defect and the protein cannot traffic past the ER and is degraded via the proteasomal degradation pathway (40). It has been suggested that FKBP38 may play a dual role during these events by negatively regulating CFTR synthesis and positively regulating ER-associated maturation of wild type and mutated form of CFTR (41). The latter functionality is PPIase-dependent (42) and FKBP38 was shown to own a hidden, calmodulin-stimulated PPIase activity (37,43,44).

Functional FKBP-ABC interaction seems not to be limited to ABCCs as judged from the fact that the activity of murine ABCB4/MDR3 transporter expressed in *S. cerevisiae* requires the presence of yeast FKBP12 (45).

Interestingly, yeast FKBP12 seems to compete with TWD1 for regulation of Arabidopsis ABCB1 activity in yeast (26,46), suggesting that this mode of action might be evolutionary conserved over kingdoms. This is further supported by the finding that FKBP12 partially complements the *twd1* mutant phenotype (26).

Here in an attempt to explore the relevance of putative proline residues that might be a substrate of a yet to be demonstrated PPIase activity of TWD1, we mapped putative surface prolines on the NBD2 of ABCB1 and tested their impact on ABCB1-mediated auxin transport activities. An essential D/E-P1.008 motif was identified that seems to be conserved in all so far described ABCBs that transport auxin and thus seems to define ATAs. The D/E-P1.008 motif is important for ABCB1-TWD1 interaction and for TWD1-mediated activation of ABCB1 transport activity supporting a previously suggested scenario in that TWD1 acts as a positive modulator of ABCB1 transport activity by means of its putative PPIase activity.

## Experimental Procedures

### Auxin transport measurements

Simultaneous ^3^H-IAA and ^14^C-benzoic acid (BA) or ^14^C-malic acid (malate) export from tobacco (*N. benthamiana*) mesophyll protoplasts was analyzed as described (26). Tobacco mesophyll protoplasts were prepared 4 days after agrobacterium-mediated transfection of Wt and indicated mutated versions of *35S:ABCB1-YFP* (26) or *35S:ABCB14-GFP* (23). In some cases, *35S:ABCB1-YFP* was co-expressed with *35S:TWD1-mCherry*. Relative export from protoplasts is calculated from exported radioactivity into the supernatant as follows: (radioactivity in the protoplasts at time t = x min.) - (radioactivity in the supernatant at time t = 0)) * (100%)/ (radioactivity in the supernatant at t = 0 min.); presented are mean values from >4 independent transfections.

### Arabidopisis thaliana

ecotype Col Wt or *abcb1-1/pgp1-1* (in Was Wt) was transformed by floral dipping with Wt and mutant versions of *35S:ABCB1-YFP* (26) or *ABCB1:ABCB1-GFP* (47), respectively. Homozygous lines were isolated by progeny analyses and used for simultaneous ^3^H-IAA and ^14^C-BA export from mesophyll protoplasts as described in (26).

### Site-directed mutagenesis

Point mutations in *ABCB1* were introduced into *35S:ABCB1-YFP* (26) and *35S:ABCB14-GFP* (23) using the QuikChange Lightning Site-Directed Mutagenesis Kit (Agilent).

### ABCB1-TWD1 interaction analyses

ABCB1 co-immunoprecipitation analyses using anti-GFP micro beads (Miltenyi Biotec, Germany) were carried using microsomal fractions prepared from Agrobacterium-co-infiltrated *N. benthamiana* in triplicates as described recently (48).

For BRET analysis, *N. benthamiana* leaves were Agrobacterium co-infiltrated with indicated BRET construct combinations (or corresponding empty vector controls) and microsomal fractions were prepared 4 d after inoculation. BRET signals were recorded from microsomes (each ∼10 µg) in the presence of 5 µM coelenterazine (Biotium Corp.) using the Cytation 5 image reader (BioTek Instruments), and BRET ratios were calculated as described previously (31). The results are the average of 20 readings collected every 30 seconds, presented as average values from a minimum of three independent experiments (independent Agrobacterium infiltrations) each with four technical replicates.

### Confocal laser scanning imaging

For confocal laser scanning microscopy work, a SP5 confocal laser microscope was used. Confocal settings were set to record the emission of GFP (excitation 488 nm, emission 500–550 nm), YFP (excitation 514nm, emission 524-550 nm), mCherry (excitation 587nm, emission 550-650nm) and FM4-64 (excitation 543 nm, emission 580-640 nm).

### Structure, sequence and phylogenetic analyses

The homology-modelled structure for Arabidopsis ABCB1 (GenBank: NP_181228) recently modelled on the high-resolution P-glycoprotein ABCB1 structure (3G5U) from *Mus musculus* (MmABCB1) was utilized as a template throughout this study (49). Human CFTR/ABCC7 (GenBank: NP_000483.3) was modeled on zebrafish CFTR (5TSI) using iTasser (https://zhanglab.ccmb.med.umich.edu/I-TASSER) because human CFTR did not contain structural information for the position relevant for D/E-P1.008. All structure figures were prepared and displayed using UCSF Chimera (https://www.cgl.ucsf.edu/chimera).

A phylogenetic analysis of Arabidopsis full-size ABCB transporter family members was performed by constructing maximum likelihood trees using PhyML 3.0 (http://www.atgc-montpellier.fr/phyml) based on the amino acid sequences aligned using MUSCLE (https://www.ebi.ac.uk/Tools/msa/muscle/). The graphical representation of the patterns within a multiple sequence alignment of selected α helices were done with WebLogo 3 (http://weblogo.threeplusone.com).

### Measurement of ATPase activity

Vanadate-sensitive ATPase activity of Wt and mutant ABCB1 was measured from microsomes (0.06 µg/ µl) prepared from transfected tobacco plants using the colorimetric determination of orthophosphate released from ATP as described previously (50). Briefly, microsomes were added to ATPase buffer (20mM MOPs, 8mM MgSO_4_, 50mM KNO_3_, 5mM NaN_3_, 0.25mM Na_2_MoO_4_, 2.5mM Phosphoenolpyruvate, 0.1% Pyruvate kinase) in the presence and absence of 0.5 mM sodium *ortho*-vanadate. The reaction was started by the addition of 15 mM ATP and incubated at 37°C for 15 min and 30 min with shaking. The stopping/ascorbate mixture was added to each well to terminate the reaction and after 10 min, the reaction mix was transferred into a stabilizing solution and incubated at RT for 1 hour. The amount of inorganic phosphate released was quantified with a colorimetric reaction, using a Cytation 5 reader (BioTek Instruments).

### Data Analysis

Transport data were statistically analyzed using Prism 7.0a (GraphPad Software, San Diego, CA).

## Results

### Identification of a conserved D/E-P motif in the C-terminal nucleotide-binding fold of ABCBs

Despite the fact that all attempts to demonstrate a PPIase activity on TWD1 were unsuccessful (36), we aimed in this study to test the impact of relevant proline residues on ABCB1 transport activity by using site-directed mutagenesis. In order to narrow down the number of candidate prolines, we focused on prolines that were potentially sterically accessible by TWD1 based on experimentally verified TWD1-ABCB1 contact sites. Yeast two-hybrid (36) and bioluminescence resonance energy transfer (BRET) (31,46) analyses allowed to limit the interacting site on ABCB1 to aa residues 980 to 1.286 of the C-terminal nucleotide binding domain. Further, by using a valid structural model for Arabidopsis ABCB1 that was built on mouse ABCB1/PGP as a template (49), we were able to cut down the numbers of putative surface-exposed prolines to the four prolines, P1.008, P1.034, P1.149 and P1.252 (Fig. 1). For doing so, we used the in-ward open conformation of ABCB1 (Fig. 1A) because we assumed that any activation of ABCB1 transport by TWD1 (26) should be occurring in this state. Finally, we included in our mutagenesis approach the two linker prolines, P635 and P647 (Fig. 1A), because ABCB linkers are thought to be very flexible and thus most-likely be also accessible by TWD1 (26,51). Another rationale was that S634 was recently shown to be phosphorylated by the AGC kinase PINOID in an action that was dependent on TWD1, resulting in altered ABCB1 transport activity (26). Interestingly, despite the tremendous re-arrangements known to occur between inward and outward-facing ABCB conformations, all prolines (except P647) remained surface-exposed also in the outward-facing conformation (Fig. 1B).

Web-based tools, such as CISPEPpred (http://sunflower.kuicr.kyoto-u.ac.jp/~sjn/cispep), predicted that all selected prolines except P1034 would be in *trans* (not shown). A thorough manual investigation of the selected 6 proline residues using both inward-open and out-ward open modelled conformers of ABCB1 revealed that all Cα-C-N-Cα bonds were actually in the *trans* conformations independent of the ABCB1 conformation (Fig. 1C). Sequence alignments with all 29 Arabidopsis ABCB isoforms indicated that the 6 prolines showed a variable degree of conservation: while the linker prolines and the very C-terminal P1252 showed a very low degree of conservation, the remaining three prolines that are part of the NBD2 were clearly more conserved (Fig. 1D, Suppl. Fig. 2). A comparison between the proline-containing stretches of NBD2 with those of NBD1, known to evolve from each other by gene duplication (52), indicated that P1.008 but not the other three prolines seemed to be specific for NBD2 (Suppl. Fig. 3).

Interestingly, we found that the presence of a D/E-P motif in the NBD2 correlated perfectly with so far diagnosed auxin transport capacities of investigated Arabidopsis ABCBs (Suppl. Figs. 1). Intriguingly, sorghum DWARF3/ABCB1 (53), maize BRACHYTIC2/ABCB1 (53), rice ABCB14 (22) and tomato ABCB4 (21), all recently demonstrated to function as auxin transporters, also contain such a conserved D/E-P motif. In summary, it appears that auxin-transporting ABCBs contain a surface-exposed, conserved D/E-P motif in the NBD2.

### Mutagenesis of the D/E-P1008 motif destroys the auxin-transport activity of ABCBs without interfering with its ATPase activity or plasma membrane presence

Introducing single point mutations in the NBDs of ABC transporters can have strong effects on protein stability most likely because non-functional ABC transporters are recognized as such and degraded (54). In order to exclude an unwanted impact of these 6 proline mutations on ABCB1 secretion and/ or protein stability, we semi-quantitatively analyzed their expression by agrobacterium-mediated leaf transfection of the tobacco, *N. benthamiana*. Confocal analyses of epidermal layers of leaves (Fig. 2A) and protoplasts prepared from those (Suppl. Fig. 4) counterstained with the PM styryl dye, FM4-64, revealed that neutralization of the 6 prolines by glycine, allowing the peptide backbone to freely rotate, did not significantly alter ABCB1-YFP expression and location. These imaging data were confirmed by quantification of Western blots on microsomal fractions from three independent leaf infiltrations (Fig. 2B) revealing similar expression levels based on ratios of ABCB1-YFP and the PM marker, PIP2;1 (26).

**Figure 2:**
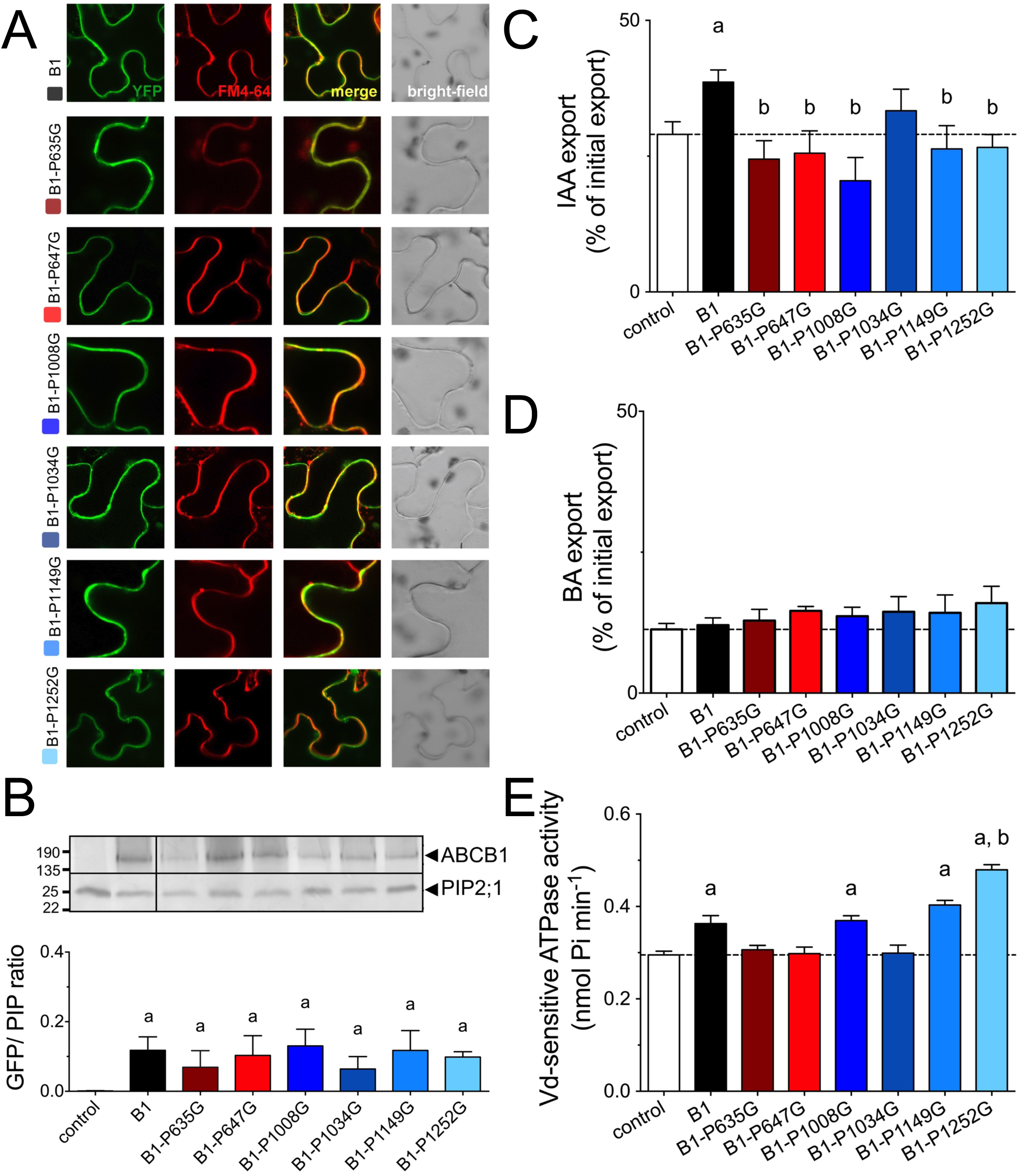
Mutation of most surface-exposed prolines in ABCB1 reduces IAA export. **(A)**Mutation of six surface-exposed prolines does not significantly alter expression and plasma membrane (PM) as revealed by confocal imaging of ABCB1-GFP of tobacco leaves transfected with ABCB1-GFP and stained with PM marker, FM4-64. **(B)**Proline mutation of ABCB1 does not significantly alter expression levels revealed by Western analyses of total microsomes prepared from tobacco leaves transfected with ABCB1 (upper panel). Microsomes were stained for GFP and plasma membrane marker, PIP2;1; ABCB1 expression was evaluated by calculating GFP/PIP ratios from 3 independent tobacco transfections (lower panel). **(C-D)** IAA (**A**) and benzoic acid (BA, **B**) export of protoplasts prepared from tobacco leaves transfected with Wt and indicated proline mutants of ABCB1. **(E)** Vanadate (Vd)-sensitive ATPase activity of microsomal fractions prepared from tobacco leaves transfected with Wt and indicated proline mutants of ABCB1; controls are in Suppl. Fig. 6 Significant differences (unpaired *t* test with Welch’s correction, p < 0.05) to vector control or ABCB1 are indicated by an ‘a’ or a ‘b’, respectively, (mean ± SE; n ≥ 4 transport experiments generated from independent tobacco transfections).

Next, we quantified IAA export of mutated version of ABCB1 from protoplasts prepared from leaf-transfected tobacco (Suppl. Fig. 4). Interestingly, except the P1.034 mutation all proline mutations revealed reduced IAA export to vector control level (Fig. 2C). This inhibitory effect was specific for IAA because transport of the diffusion control, benzoic acid (BA), was not affected by ABCB1 mutation (Fig. 2D). In order to test if proline mutations of the NBD2 and eventually also the linker had an effect on ATP hydrolysis by ABCB1, we quantified ATPase activity of microsomal fractions prepared from transfected tobacco. ABCB1 expression significantly increased the fraction of vanadate-sensitive ATPase activity (Fig. 2E), which can be attributed to ABC transporters, while no significant difference between vector control and ABCB1 was found for the vanadate-insensitive fraction or in the absence of ATP (Suppl. Fig. 5). ABCB1-mediated ATPase activity was not stimulated by the native substrate, IAA, as found for many ABC transporters (55). Interestingly, loss of IAA transport for both linker mutations correlated with a loss of ATPase activity, supporting the proposed impact of the regulatory linker on the NBD motions (56). The very C-terminal proline mutation showed increased ATPase activity, which indicates that this mutation might have resulted in an uncoupling of transport from ATPase activity. The most interesting finding, however, was that P1.008G mutation did abolish IAA transport without affecting ABCB1 ATPase activity, indicating that P1.008 most likely interferes with ABCB1 activity regulation by a mode that is essentially independent of ATP hydrolysis.

In light of the proposed, essential function of the D/E-P1.008 motif for auxin transport, we asked if the D/E prior to the proline was likewise critical. Interestingly, alanine substitution of E1.007 also reduced IAA transport but not BA transport activity (Fig. 3A-B) without changing significantly ABCB1 expression or location (Fig. 3D). Similarly, E1.007A mutation had no impact on ABCB1 ATPase activity (Fig. 3C), indicating in summary that both residues of the D/E-P1.008 motif are essential for ABCB1 IAA transport in a mode that most likely does not involve the ATPase activity of NBD2.

**Figure 3:**
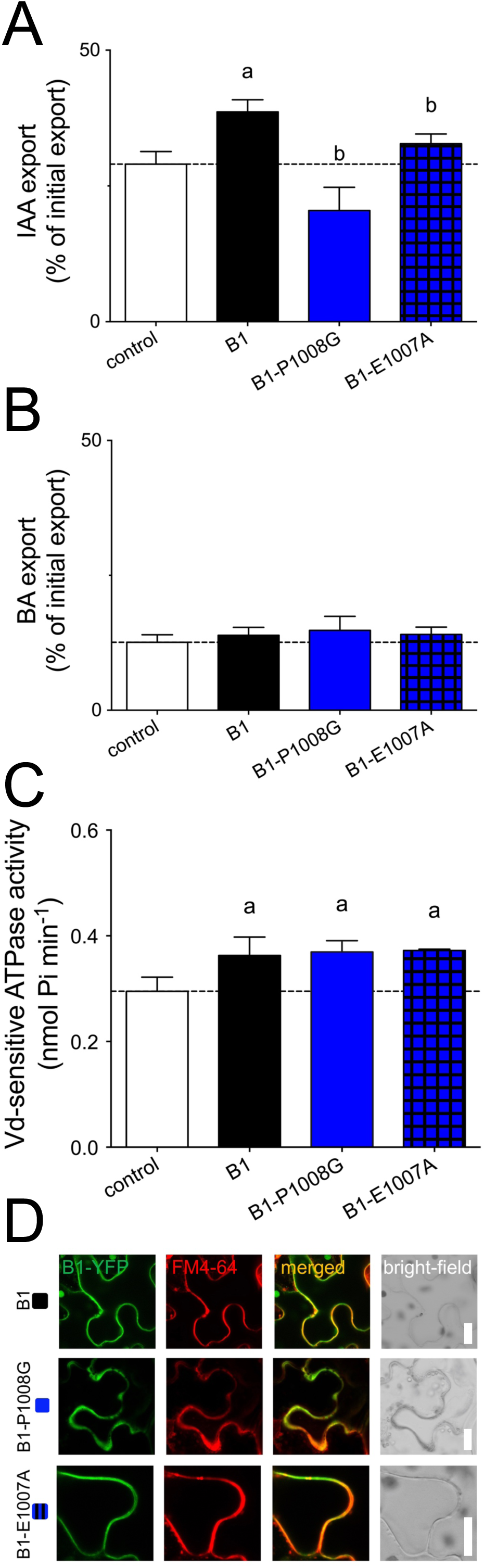
Mutagenesis of the D/E-P1008 motif abolishes the auxin-transport activity of ABCB1 without interfering with its ATPase activity or PM location. **(A-B)** IAA (**A**) and benzoic acid (BA, **B**) export of protoplasts prepared from tobacco leaves transfected with Wt and indicated D/E-P1.008 mutations of ABCB1. **(C)**ATPase activity of microsomal fractions prepared from tobacco leaves transfected with Wt and indicated D/E-P1008 mutations of ABCB1. Significant differences (unpaired *t* test with Welch’s correction, p < 0.05) to vector control or ABCB1 are indicated by an ‘a’ or a ‘b’, respectively, (mean ± SE; n ≥ 4 transport experiments generated from independent tobacco transfections). **(D)** D/E-P1.008 mutants of ABCB1 are expressed at similar level on the plasma membrane revealed by confocal imaging of ABCB1-GFP of tobacco leaves transfected with ABCB1-GFP and stained with PM marker, FM4-64.

### Introduction of a D/E-P motif into malate importer, ABCB14, increases its background auxin transport activity

The only Arabidopsis ABCB that does not transport auxin but for that a transport activity has been proven is ABCB14 functioning as a PM malate importer (23). ABCB14 contains P-P991 instead of the D/E-P. PM expression of Wt ABCB14 in tobacco resulted in slightly reduced malate export (an indicative for enhanced import) verifying previous studies (23) but also a slightly increased IAA export, although both differences were not significantly different from vector control (Fig. 4A-B). Interestingly, correction of the P-P991 into E-P991 both significantly enhanced malate import by ABCB14, as well as IAA export activity without changing its PM location (Fig. 4C). In summary, this data set indicates that introduction of a D/E-P motif into a non-auxin transporting ABCB increases its native (malate) transport capacity and thus most likely also a background IAA export activity that was suggested recently (17).

**Figure 4:**
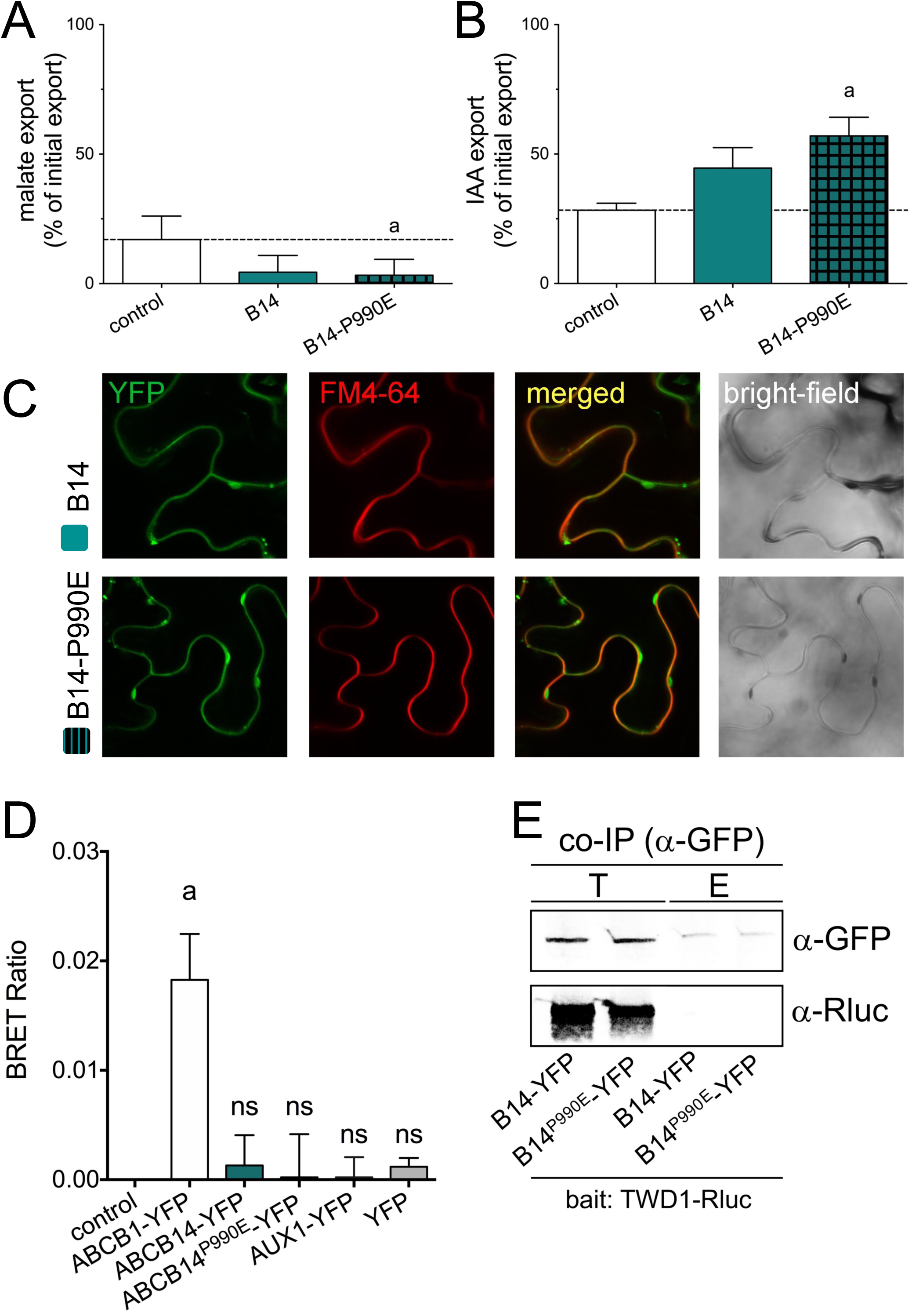
Introduction of an E-P motif into malate importer, ABCB14, increases its transport activity. **(A-B)** Simultaneous malate (**A**) and IAA (**B**) export of protoplasts prepared from tobacco leaves transfected with Wt and P990E-mutated ABCB14. **(C)**P990E mutation does not alter significantly expression or plasma membrane location of ABCB14 revealed by confocal imaging of ABCB1-GFP of tobacco leaves transfected with ABCB1-GFP and stained with PM marker, FM4-64. **(D)** BRET analyses of microsomal fractions prepared from tobacco leaves co-transfected with TWD1-Rluc and Wt and P990E mutation of ABCB14-GFP. TWD1-Rluc alone, AUX1-YFP/TWD1-Rluc and YFP/TWD1-Rluc were used as negative and ABCB1YFP/TWD1-Rluc as positive controls. Significant differences (unpaired *t* test with Welch’s correction, p < 0.05) to ABCB1-YFP/TWD1-Rluc are indicated by an ‘a’ (mean ± SE; n=3 BRET measurements from 3 independent tobacco transfections). **(E)**Co-immunoprecipitation of TWD1-Rluc using Wt and P990E mutation of ABCB14-YFP as a bait. Co-IP was performed from microsomal fractions prepared from tobacco leaves co-transfected with indicated combinations. Total input (T) and elution (E) were probed against anti-GFP and anti-Rluc detecting ABCB14-YFP and TWD1-Rluc, respectively.

### The D/E-P1.008 motif is important for ABCB1 interaction with TWD1

The findings above left us with the possibility that the D/E-P1.008 motif in ABCB1 was not only essential for ABCB1 activity regulation but eventually also for interaction with TWD1 described as a regulator of ABCB1 transport activity by protein-protein interaction (26,31,46,57). In fact, it was shown that dissociation of TWD1 from ABCB1 by the auxin efflux inhibitor, NPA, disrupted the regulatory impact of TWD1 on ABCB activity (46). In order to test the impact of the E-P1.008 motif on TWD1-ABCB1 interaction, we measured TWD1-ABCB1 interaction by employing an established microsome-based BRET assay (31,46). As shown before, co-expression of ABCB1-YFP and TWD1-Rluc resulted in significant BRET ratios, while the negative controls, AUX1-YFP or YFP alone, did not (Fig. 5A). Interestingly, also co-expression of ABCB4-YFP with TWD1-Rluc resulted in comparable BRET ratios (Fig. 5A) without significantly altering expression or location of both partners (Supp. Fig. 6). This supports the concept that lack of ABCB4-TWD1 interaction is the cause of ABCB4 de-regulation in *twd1* (31,32).

**Figure 5:**
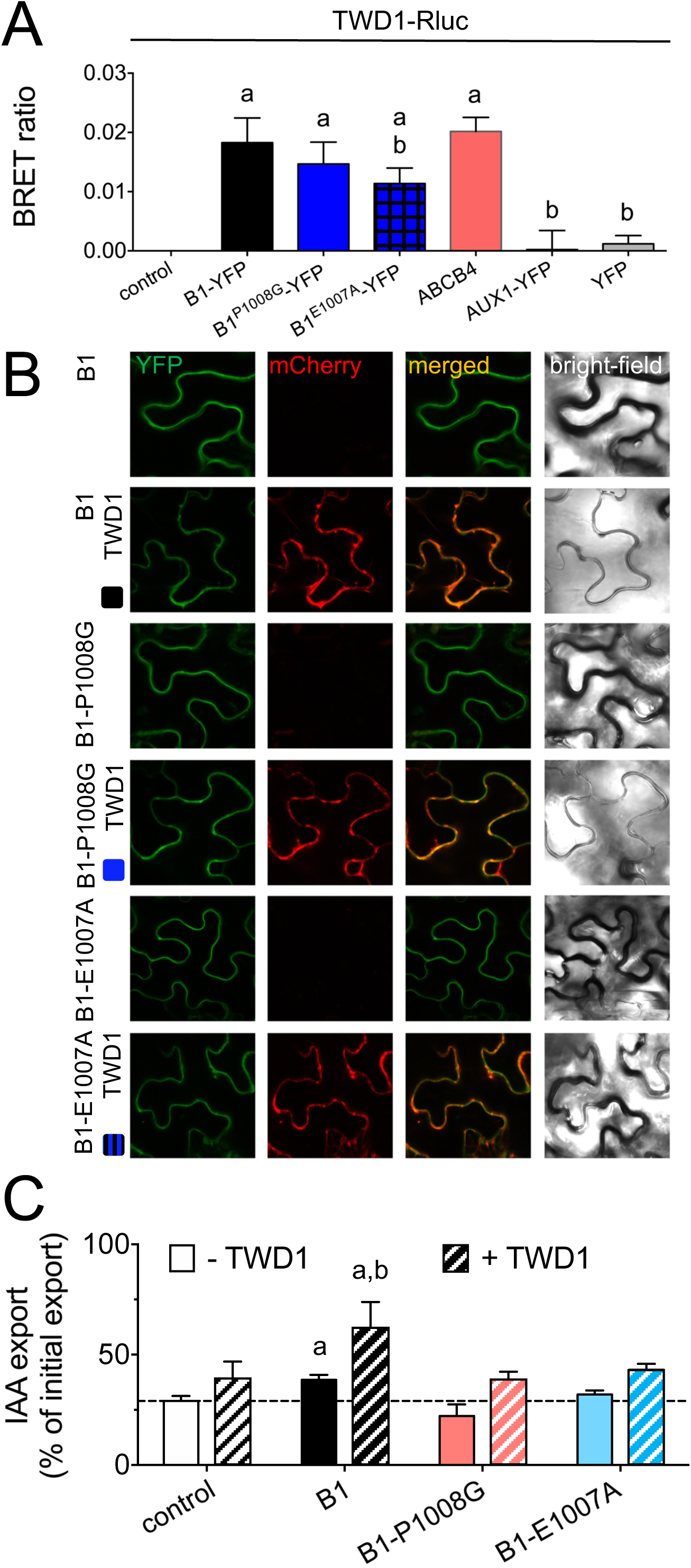
The D/E-P1008 motif in ABCB1 is important for interaction with TWD1 and regulation by TWD1. **(A)** BRET analyses of microsomal fractions prepared from tobacco leaves co-transfected with TWD1-Rluc and Wt and indicated D/E-P1.008 mutations of ABCB1-YFP. TWD1-Rluc alone, AUX1-YFP/TWD1-Rluc and YFP/TWD1-Rluc were used as negative controls. Significant differences (unpaired *t* test with Welch’s correction, p < 0.05) to TWD1-Rluc alone are indicated by an ‘a’, significant differences to ABCB1-YFP/TWD1-Rluc by a ‘b’ (mean ± SE; n=3 BRET measurements from 3 independent tobacco transfections). **(B)** Both E1.007A and P1.008G mutant versions of ABCB1-YFP co-localize with TWD1-mCherry on the epidermal PM of tobacco leaves after co-transfection. **(C)**IAA export of protoplasts prepared from tobacco leaves co-transfected with TWD1-mCherry and Wt or indicated D/E-P1.008 mutations of ABCB1-YFP. Significant differences (unpaired *t* test with Welch’s correction, p < 0.05) to vector control or ABCB1-YFP are indicated by an ‘a’ or a ‘b’, respectively (mean ± SE; n=4 transport measurements from 4 independent tobacco transfections).

**Figure 6:**
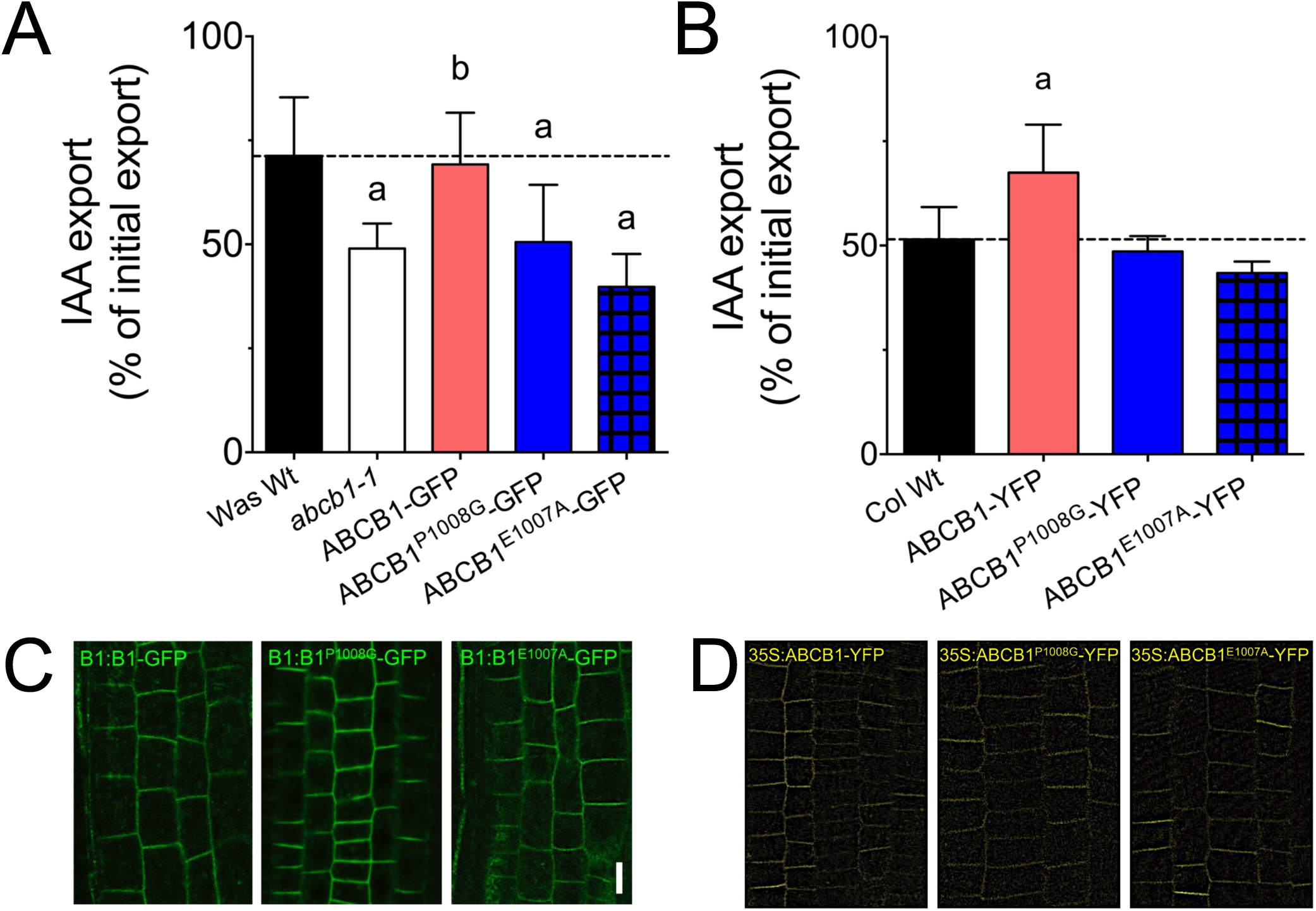
The D/E-P1.008 motif in ABCB1 is essential for auxin transport activity *in planta*. **(A)** IAA export of protoplasts prepared from *abcb1-1* transformed with Wt or indicated E-P1.008 mutations of ABCB1-GFP expressed under its native promoter (ABCB1:ABCB1-GFP). **(B)** IAA export of protoplasts prepared from Col Wt transformed with Wt or indicated E-P1.008 mutations of ABCB1YFP expressed under a constitutive promoter (35S:ABCB1-YFP). Significant differences (unpaired *t* test with Welch’s correction, p < 0.05) to ABCB1-YFP/TWD1-Rluc are indicated by an ‘a’ (mean ± SE; n=3 BRET measurements from 3 independent tobacco transfections). Benzoic acid diffusion controls are in Suppl. Fig. 10 (**C-D**) Confocal analyses of Wt and mutant versions of ABCB1:ABCB1-GFP in *abcb1-1* (**C**) or 35S:ABCB1-YFP in Col Wt (**D**), respectively, revealed no significant differences in BCB1 expression and location.

Excitingly, both mutation of P1.008G (ca. 17% reduction) and E1.007A in ABCB1-YFP (ca. 39% reduction) decreased interaction with TWD1-Rluc, while only the latter was significant (Fig. 5A). Co-expression of TWD1 with mutant version of ABCB1 did not significantly alter expression or PM location of both partners excluding unwanted secondary effects (Fig. 5B). In summary, these data indicate that the E-P.1008 motif is important but not essential for ABCB1-TWD1 interaction.

### The E-P1.008 motif in ABCB1 is essential for ABCB1 activation by TWD1 and auxin transport activity *in planta*

In order to evidence that the described activation of ABCB1-mediated IAA export caused by TWD1 interaction is dependent on the E-P1.008 motif, we quantified IAA export from protoplasts prepared from tobacco leaves co-infiltrated with TWD1-mCherry and Wt and mutant versions of ABCB1-YFP. As described before (26), TWD1-mCherry enhanced slightly IAA export of vector controls, most likely due to activation of tobacco-endogenous ABCBs (Fig. 5C). In contrast, co-transfection of TWD1-mCherry with Wt ABCB1-YFP significantly enhanced IAA export. Strikingly, the activating effect of TWD1-mCherry was absent with E1.007 and P1.008 mutations of ABCB1-YFP clearly indicating that the E-P1.008 motif in ABCB1 is essential for the regulatory impact provided by TWD1.

Finally, in order to test the importance of the E-P1008 motif *in planta*, we expressed Wt and E1.007A and P1.008G mutant versions of ABCB1 in *abcb1-1* and Col Wt under its native promoter (ABCB1:ABCB1-GFP) and a constitutive one (35S:ABCB1-YFP), respectively. As expected, complementation of the *abcb1-1* mutant with WT but not both mutant versions rescued IAA efflux to Was Wt levels (Fig. 6A). In agreement, constitutive over-expression in Col Wt revealed enhanced IAA export in comparison to Wt only for Wt ABCB1 but not for both mutants. Like in tobacco no differences were found with the diffusion control, benzoic acid (Suppl. Fig. 7). Importantly, confocal analyses of mutant versions of ABCB1:ABCB1-GFP in *abcb1-1* (Fig. 6C) and 35S:ABCB1-YFP in Col Wt (Fig. 6D), respectively, revealed no significant differences in ABCB1 expression and location in comparison to Wt ABCB1. In summary, this data set underlines the notion that the D/E-P1.008 motif in auxin-transporting ABCBs is essential for auxin transport activity *in planta*.

## Discussion

### A conserved D/E-P motif in the C-terminal nucleotide binding domain defines auxin-transporting ABCBs (ATAs)

By using a structure-based approach we have identified a series of surface-embedded proline residues in the NBD2 and linker of Arabidopsis ABCB1 (Fig. 1) that do not alter ABCB1 protein stability or location but its catalytic transport activity (Figs. 2-3). Of special interest was P1.008 because we uncovered that this proline is part of a signature D/E-P motif that seems to be specific for ATAs. Beside the proline, also mutation of the acidic moiety prior to the proline abolishes auxin transport activity by ABCB1 in the heterologous tobacco system (Fig. 3) as well as *in planta* (Fig. 6). So far, all Arabidopsis ABCBs for that auxin transport was shown carry this conserved motif underlining its diagnostic potential. Interestingly, this motif does not seem to be restricted to dicots because it was also found in the monocot ATAs, sorghum DWARF3/ABCB1 (53), maize BRACHYTIC2/ABCB1 (53), and rice ABCB14 (22). The D/E-P signature was further found in all five ABCBs of the Arabidopsis ABCB15-22 cluster (Suppl. Fig. 1) for that no auxin transport data have yet been published. As a proof of concept, we verified specific IAA transport for all five ABCBs (Hao, Vanneste, Shani and Geisler and Beekman, unpublished data), indicating that most likely 11 out of 22 full-size Arabidopsis ABCBs do transport auxin.

To further validate our assumption, we corrected a P-P991 motif in Arabidopsis ABCB14 that was previously proposed (beside ABCB15) to promote auxin transport since inflorescence stems in both mutants showed a reduction in polar auxin transport (17). Mutation of P990E significantly increased a background auxin export activity on ABCB14 (Fig. 4), indicating that this mutation might have an impact on substrate specificity. This concept is indirectly supported by NBD mutations in the yeast ABCG-type transporter, PDR5, that altered beside transport capacity also substrate specificity without changing the ATPase activity (58). A more likely option, however, is that mutation of this motif enhances the transport capacity in general, which is supported by the fact that the P990E mutation in ABCB14 also promotes malate import (Fig. 4).

### A new mode of ABC transporter regulation by *cis-trans* isomerization of peptidyl-prolyl bonds via FKBPs?

In contrast to all other proline mutations, P1.008G and E1.007A mutations both did only affect auxin transport but not ATPase activity of ABCB1 (Figs. 2-3). A rationale for this finding is given by the fact that this motif in contrast to most other prolines is distant to the elements that are responsible for ATP binding and hydrolysis, like Walker A and B motifs (Fig. 1, Suppl. Fig. 8). Instead, E-P1.008 lies in proximity and in between intracellular/ coupling helices 2 and 3, establishing the connection between intracellular loops, ICL2 and ICL3, and the NBD2 (Fig. 1 and Suppl. Fig. 8 (59)). Mutations in intracellular/ coupling helices in human ABCB1 close to a well-conserved phenylalanine were reported to lead to dysfunctionality of the protein and its degradation (60). In Arabidopsis ABCB1, mutation of F792, part of the coupling helix 3, leads to reduced binding of the non-competitive auxin transport inhibitor, BUM, thought to bind to this region of NBD2 (61). Further, many mutations of ABCC7/ CFTR in the ICL2 and ICL4 are associated with cystic fibrosis (62).

However, an alternative but also plausible scenario is that mutation of P1.008G and/or P1.007E has a direct, steric impact on positioning of TMH12 that is connected via a short α-helix with this motif (Suppl. Fig. 8). A recent analysis of the surface residues surrounding the putative IAA binding region 3 revealed a direct involvement of TMH12 in IAA binding in analogy to human ABCB1 (49). This proposes a scenario in that TMH12 might be slightly dislocated upon a transient E-P1.008 isomerization opening the central substrate cavity. This scenario is supported by Normal Mode Analysis (NMA) of protein dynamics simulations using DynaMut (http://biosig.unimelb.edu.au/dynamut) revealing that P1.008G (but not E1.007) mutation has a significant effect on ABCB1 local rigidity (not shown). In summary, this supports the idea that structural changes triggered by proline isomerization via TWD1 in a loop close to or in the ICLs connecting TMDs with NBDs might have a momentous impact on ABCB transport activity that is absent in the E-P1.008 mutants.

Another interesting finding was that the E-P1.008 motif is not only indispensable for ABCB1 transport but also important - although not essential - for ABCB1 interaction with TWD1 (Fig. 5). Remarkably, beside the proline also mutation of the acidic moiety prior to the proline leads to reduced binding of TWD1, which might indicate that both residues are an essential part of the TWD1 docking surface on the NBD2 of ABCB1. These two lines of evidence further support a previously suggested scenario in that TWD1 acts as a positive modulator of ABCB1 transport by means of its catalytic PPIase activity (9,26,36). This activity, although not experimentally verified for TWD1, is usually associated with its PPIase domain that was shown to physically interact with a stretch of the mapped ABCB1 C-terminus (36). Strikingly, the functional human ortholog of FKBP42/TWD1, FKBP38, was demonstrated to have an impact on the biogenesis of human CFTR/ABCC7 (37). This action was shown to depend on a hidden PPIase activity on FKBP38 that was uncovered to require calmodulin activation (43). If TWD1 does also harbor such a calmodulin-stimulated PPIase activity is currently unknown but can now be tested, especially in knowledge of a putative *in vivo* peptide substrate.

Another open question is if all ATAs do physically interact with TWD1 or other, not yet identified PPIases. We here show that TWD1, beside with ABCB1,19 (12,36,46), also interacts with ABCB4 (Fig. 4), whose PM presence was shown to depend on TWD1 (31,32). Interestingly, correction of a P-P991 into an E-P991 motif in ABCB14 increased transport capacities but did not restore interaction with TWD1 shown by BRET and co-IP (Fig. 4D-E), indicating that interaction is provided by a structurally conserved surface that extends these two amino acids. However, in contrast to the described CFTR/ FKBP38 model our data suggest a mode of ABC transporter regulation in that *cis-trans* isomerization catalyzed by the PPIase activity of TWD1 would lead to a transient activation of auxin transport activity. This scenario is supported by our finding that IAA transport of Wt ABCB1 but not of E1.007A and P1.008G mutant versions of ABCB1-YFP can be activated by TWD1 (Fig. 5C). The difference and novelty of this so far not demonstrated mode of transporter regulation would lie in the fact that deactivation would not require a second enzymatic step (like for dephosphorylation) but would happen by spontaneous re-isomerization known to occur slowly (in seconds to minutes) due to the high energy barrier imposed by the partial double bond character of the peptide bond (63). Interestingly, also for HsCFTR/ ABCC7, a mutation of prolines was shown to alter its channel properties (64). Strikingly, P1.008 of AtABCB1 seems to be conserved and surface exposed also in CFTR (Suppl. Fig. 8A-B) but also in AtABCC1/2, both shown to interact with and to be regulated by TWD1 (65). Moreover, in the human Breast Cancer Resistance Protein, BCRP/ABCG2, mutation of P392, located at the TMD-NBD interface, altered transport activity and substrate specificity (66). Currently, it is unclear if these X-P bonds are also folded by FKBPs; if so these data would suggest that this mode of transporter regulation is more frequent than thought.

## Supporting information

Suppl. Infomation

## Funding and additional information

We would like thank L. Charrier for excellent technical assistance and Y. Lee for the 35S:ABCB14-GFP plasmid. This work was supported by grants from the Swiss National Funds (31003A-165877/1) and the ESA (CORA grant LIRAT) (all to MG). KP was supported by the PhD program POWR.03.02.00-00-1032/16 of Polish National Center for Research and Development.

## Author contributions statement

MG designed research; PH, KP, JL and JX performed research; PH, KP, MdD, MJ, AB and MG analyzed data; MG wrote the manuscript.

## Abbreviations

IAA: indole-3-acetic acid;
BA: benzoic acid;
PAT: polar auxin transport;
ABCB: ATP-binding cassette protein subfamily B;
PGP: P-glycoprotein;
TWD1: Twisted Dwarf1;
DGT: DIAGEOTROPICA;
MDR: multi-drug resistance;
NPA: 1-*N*-naphthylphthalamic acid;
FKBP: FK506-binding protein;
WT: wild-type;
NBD: nucleotide binding domain;
TMD: transmembrane domain;
ICL: intracellular loop;
FKBD: FK506-binding domain;
AUX1/LAX: AUXIN1-RESISTANT1/LIKE;
PIN: PIN-FORMED;
CFTR: cystic fibrosis transmembrane conductance regulator;
PPIase: *cis-trans* peptidyl-prolyl isomerase;
PM: plasma membrane;
ER: endoplasmic reticulum;
BA: benzoic acid.

