## Supplementary material for "A conserved D/E-P motif in the nucleotide binding domain of plant ABCB/PGP-type ABC transporters defines their auxin transport capacity": Suppl. Infomation

**Content:**

Supporting figures S1-S8

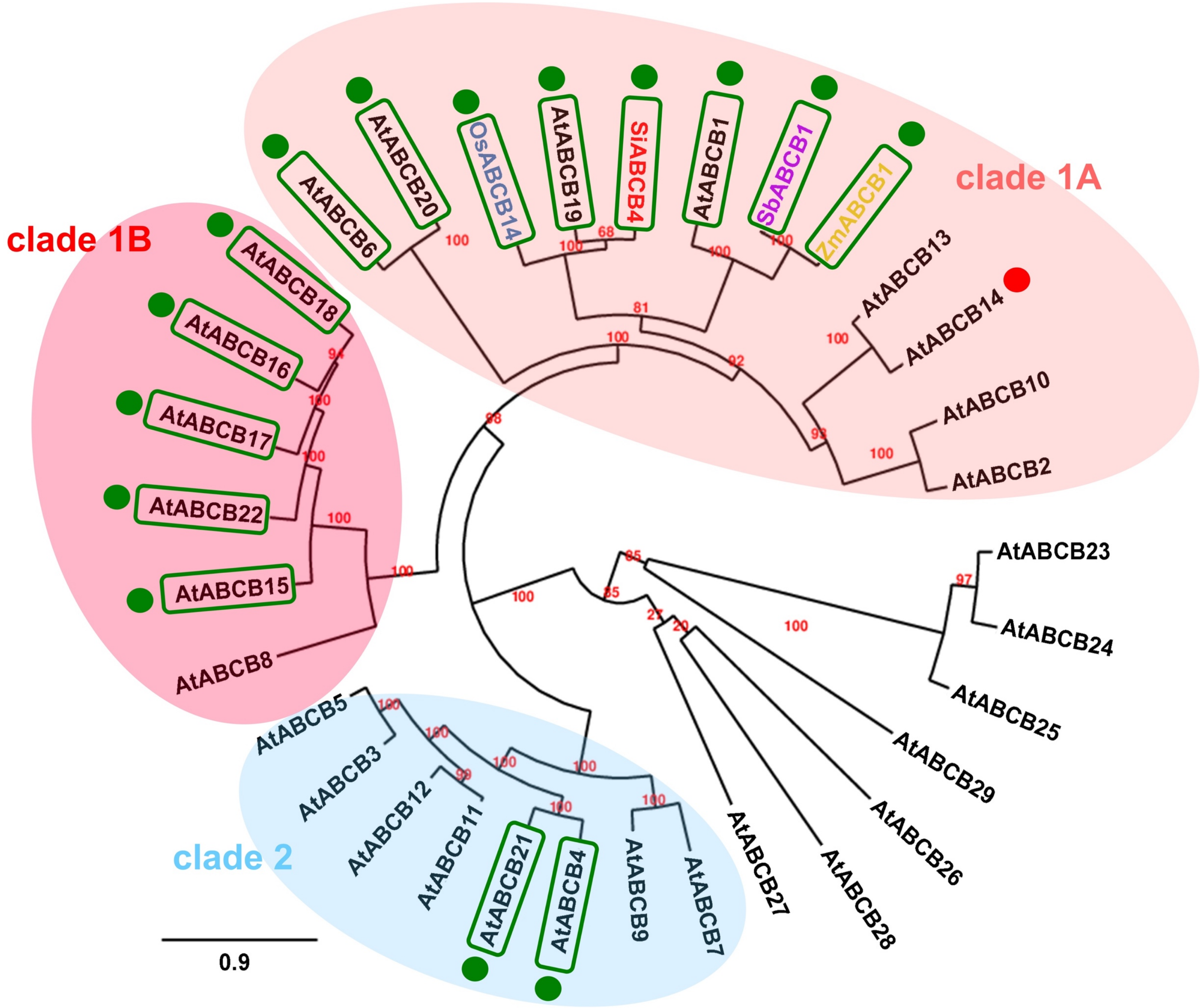


**Supporting. Fig. 1: Phylogenetic analysis of Arabidopsis ABCB transporter family members.** Phylogenetic analysis was performed by constructing maximum likelihood trees using PhyML 3.0 (http://www.atgc-montpellier.fr/phyml/) based on the amino acid sequences aligned using MUSCLE (Edgar et. al., 2004). Selected plant ABCBs from rice (Os), tomato (Si), sorghum (Sb) and maize (Zm) for that auxin transport was experimentally conformed are included. Clades of full-size ABCBs are coloured, half-size ABCBs are not. The presence of a conserved D/E-P1.008 motif (relative to AtABCB1) is indicated by green boxes; experimentally verified or falsified auxin transport is labelled with green or red dots, respectively.

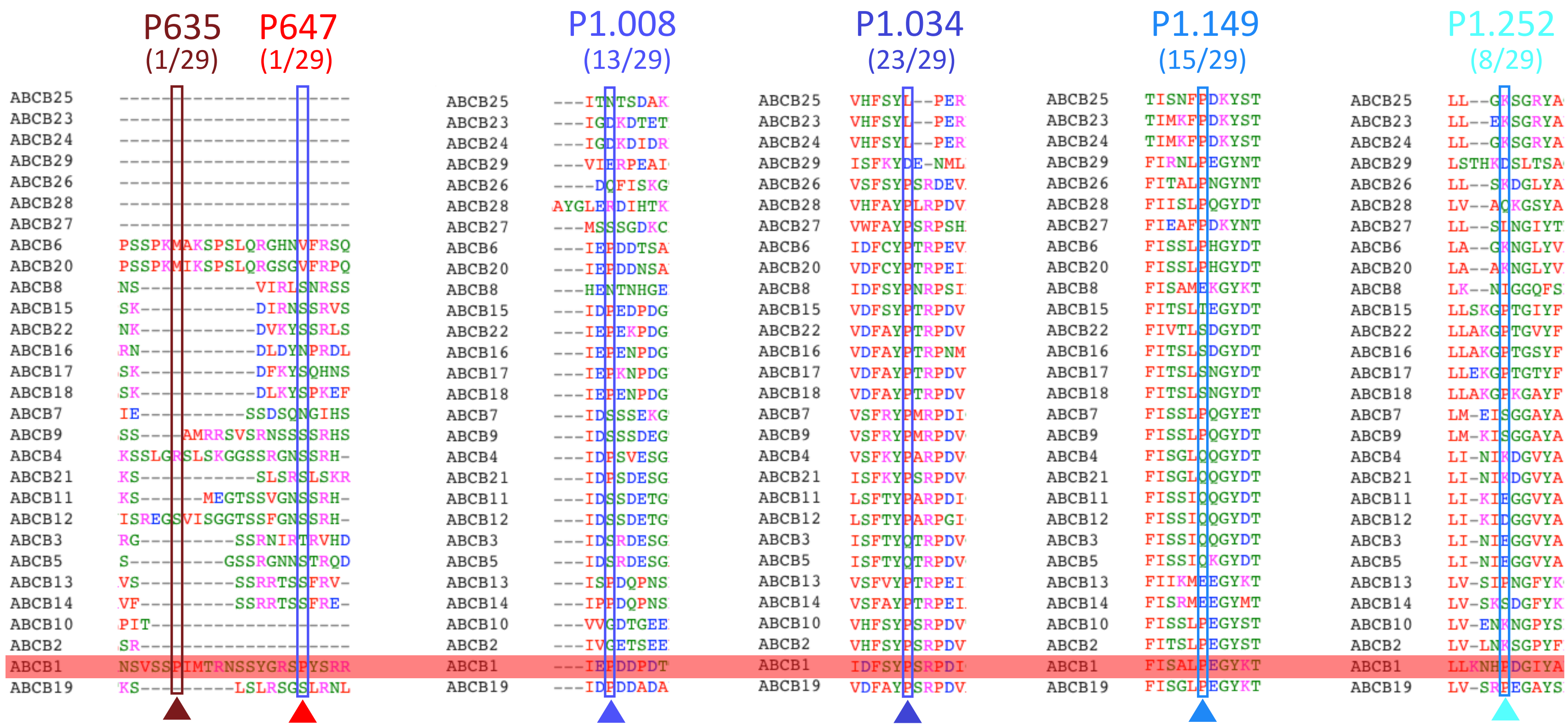


**Supporting Fig. 2: Sequence alignment of surface-exposed prolines identified in the linker and NBD2 of ABCB1 with all Arabidopsis ABCB transporter family members.** Sequences were aligned using MUSCLE (Edgar *et. al.* 2004). Conservation of prolines (indicated with triangles) amongst all 29 Arabidopsis ABCBs is indicated.

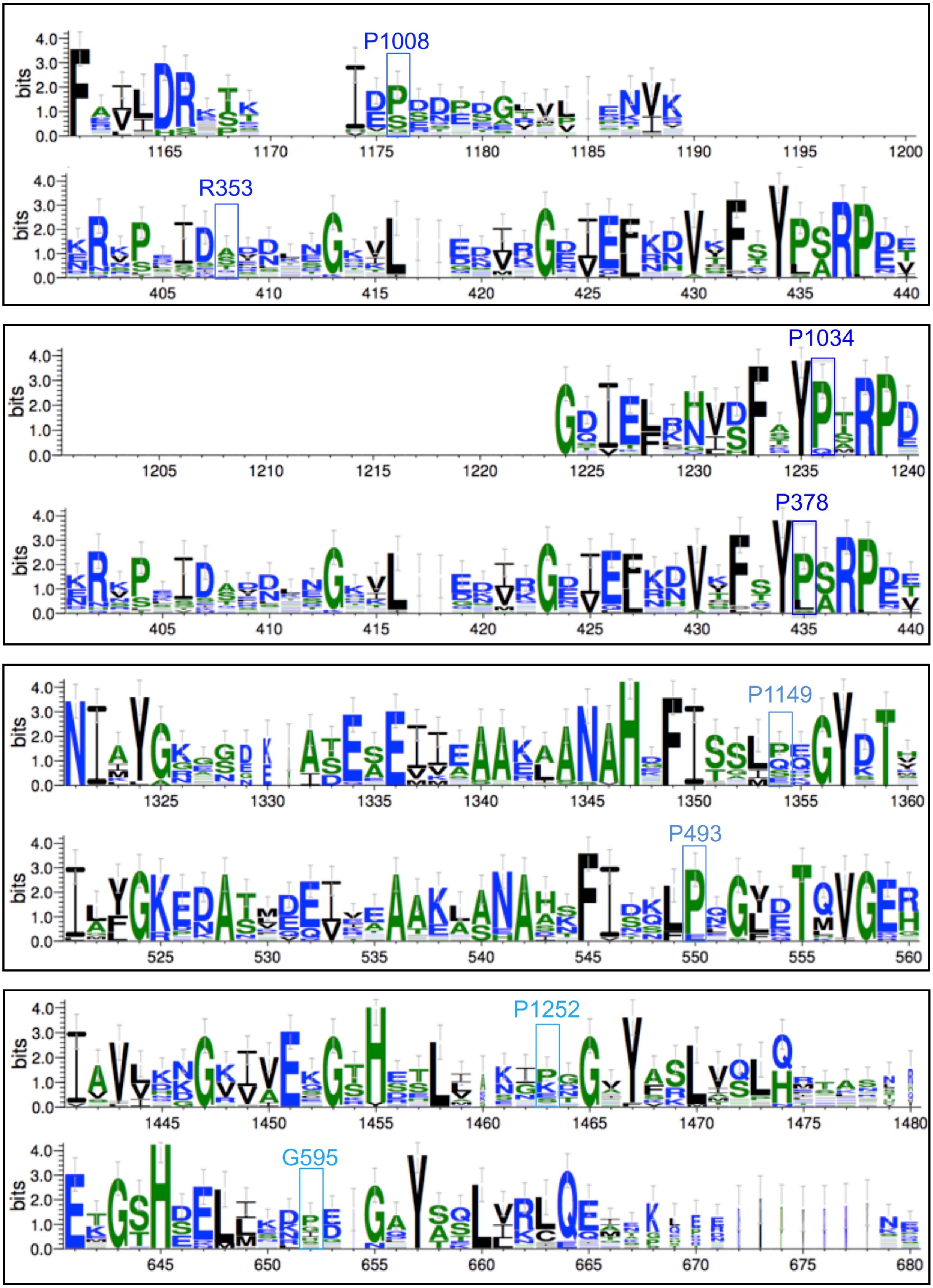


**Supporting Fig. 3: Logo representation (WebLogo 3) of NBD2 proline conservation (upper panels) in comparison with the homologous regions of NBD1 (lower panels) after multiple sequence alignment of Arabidopsis ABCBs using MUSCLE (Edgar *et. al.* 2004).**

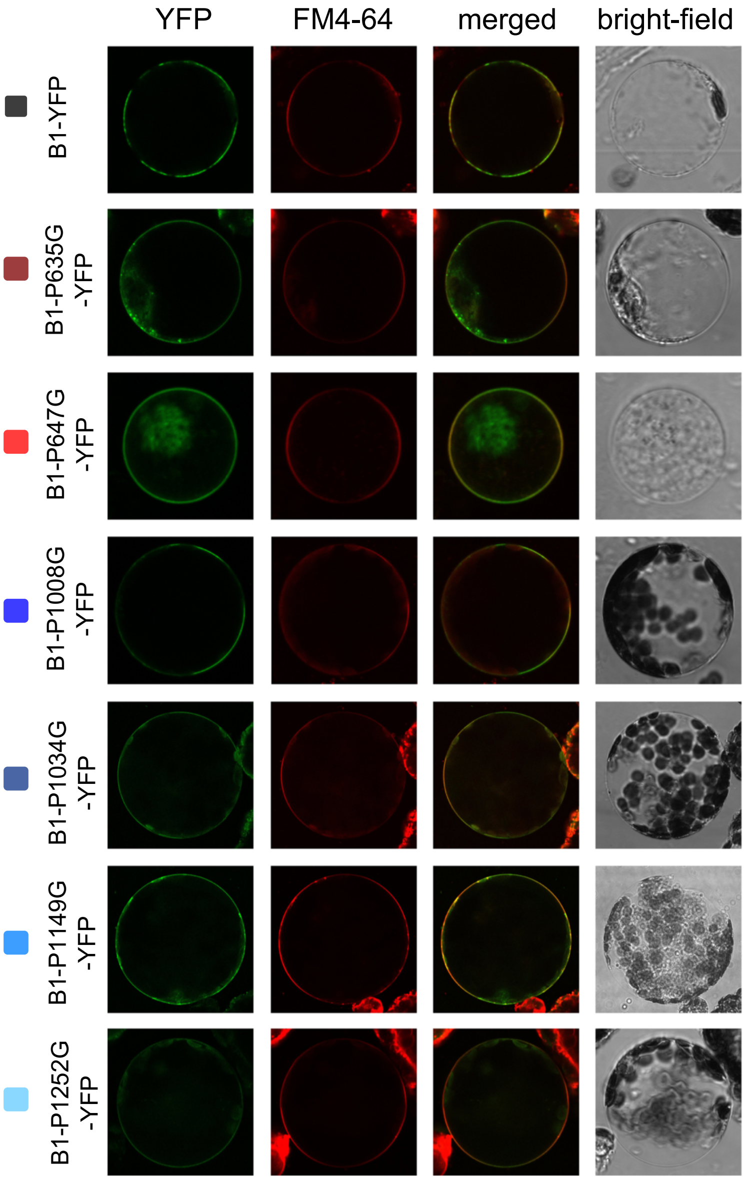


**Supporting Fig. 4: Mutation of six surface-exposed prolines in ABCB1 does not significantly alter PM location and expression on tobacco protoplasts.**

Wild-type and proline mutants of ABCB1 are expressed on the plasma membrane revealed by confocal imaging of ABCB1-GFP of tobacco protoplasts prepared from leaves transfected with ABCB1-GFP and stained with PM marker, FM4-64.

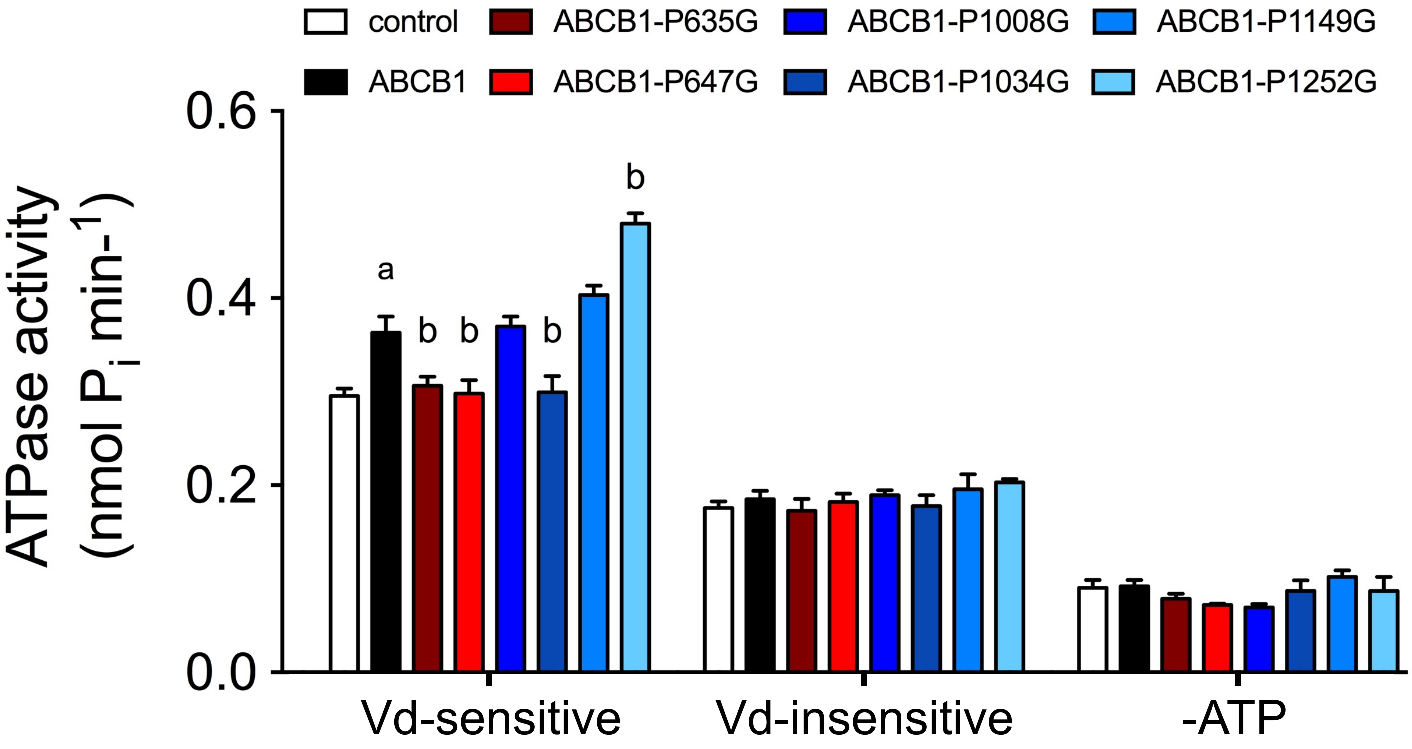


**Supporting Fig. 5: Effect of mutation of surface-exposed prolines on ATPase activity.** ATPase activity of microsomal fractions prepared from tobacco leaves transfected with wild-type and indicated proline mutants of ABCB1 measured in the presence and absence of ortho-vanadate and ATP. Significant differences (unpaired *t* test with Welch's correction, $p < 0.05$) to vector control or ABCB1 are indicated by an 'a' or a 'b', respectively, (mean ± SE; n ≥ 4 transport experiments generated from independent tobacco transfections).

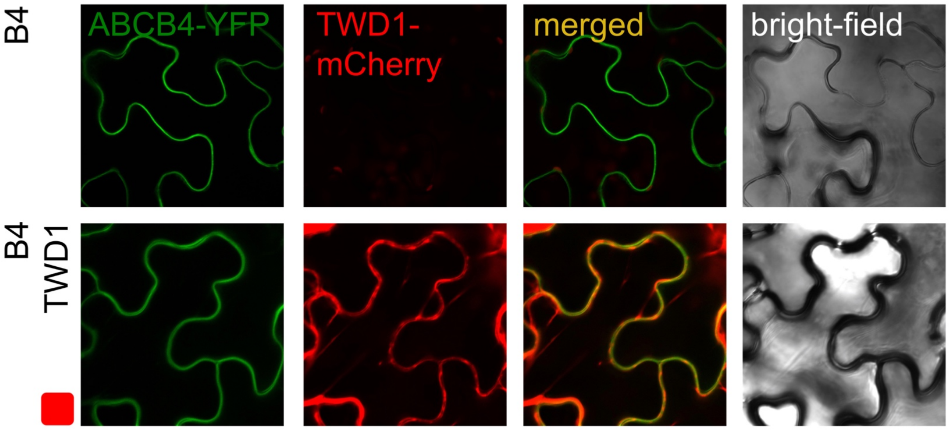


**Supporting Fig. 6: Co-expression with TWD1 does not significantly alter expression of ABCB4.**

Confocal imaging of ABCB4-GFP in the absence (upper row) and presence of TWD1-mCherry in tobacco leaves transfected with ABCB1-GFP and TWD1-mCherry.

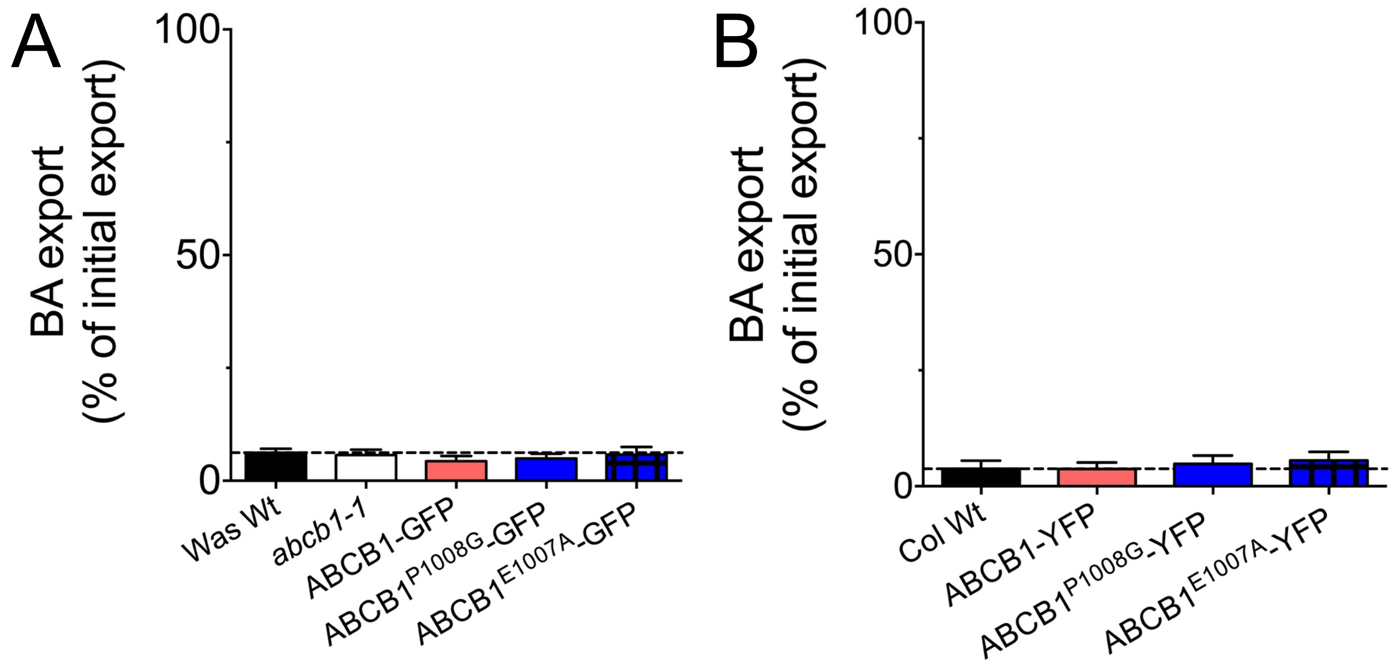


**Supporting Fig. 7: Benzoic acid (BA) export of protoplasts prepared from indicated Arabidopsis lines.** *Abcb1-1* (**A**) or Col Wt (**B**) were transformed with Wt or indicated P1.008 mutations of ABCB1 expressed under native (**A**) or constitutive promoters (**B**).

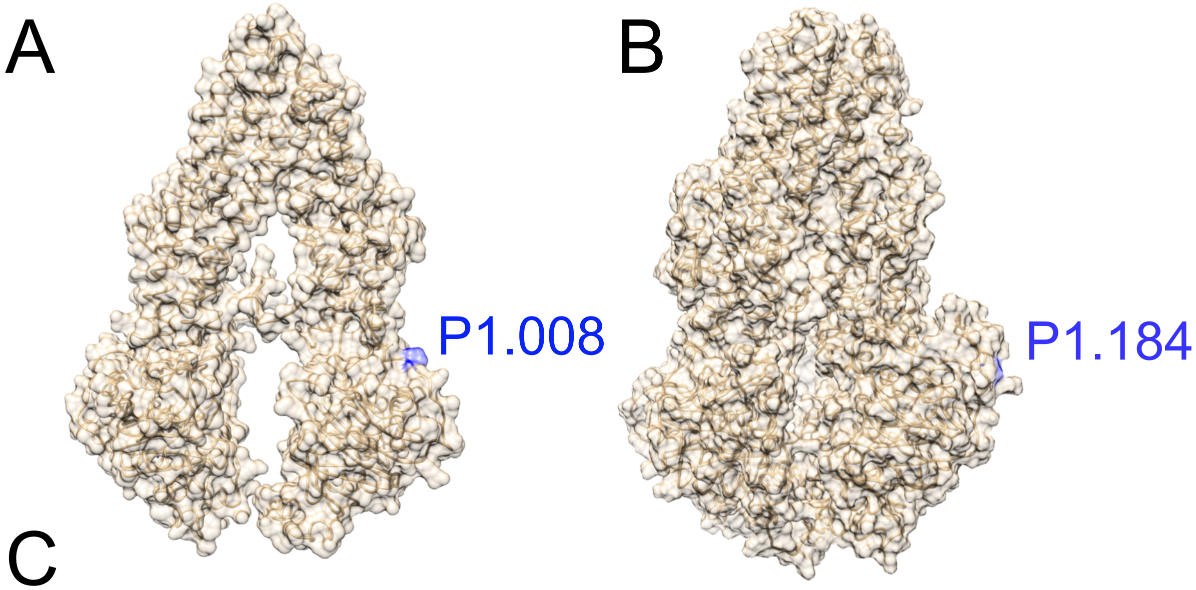


C

```
AtABCB1    IKGGQAMRSVFELLDRKTEIEP-----------------DDPDTTPVPD---RLRGEVELK
HsCFTR     DSLMRSVSRVFKFIDMPTEGKPTKSTKPYKNGQLSKVMIIENSHVKKDDIWPSGGQMTVK
AtABCC1    ENSLNSVERVGNYIEIPSE-APL----------------VIENNRPPPG--WPSSGSIKFE
AtABCC2    ENSLNAVERVGNYIEIPPE-APP----------------VIENNRPPPG--WPSSGSIKFE
```

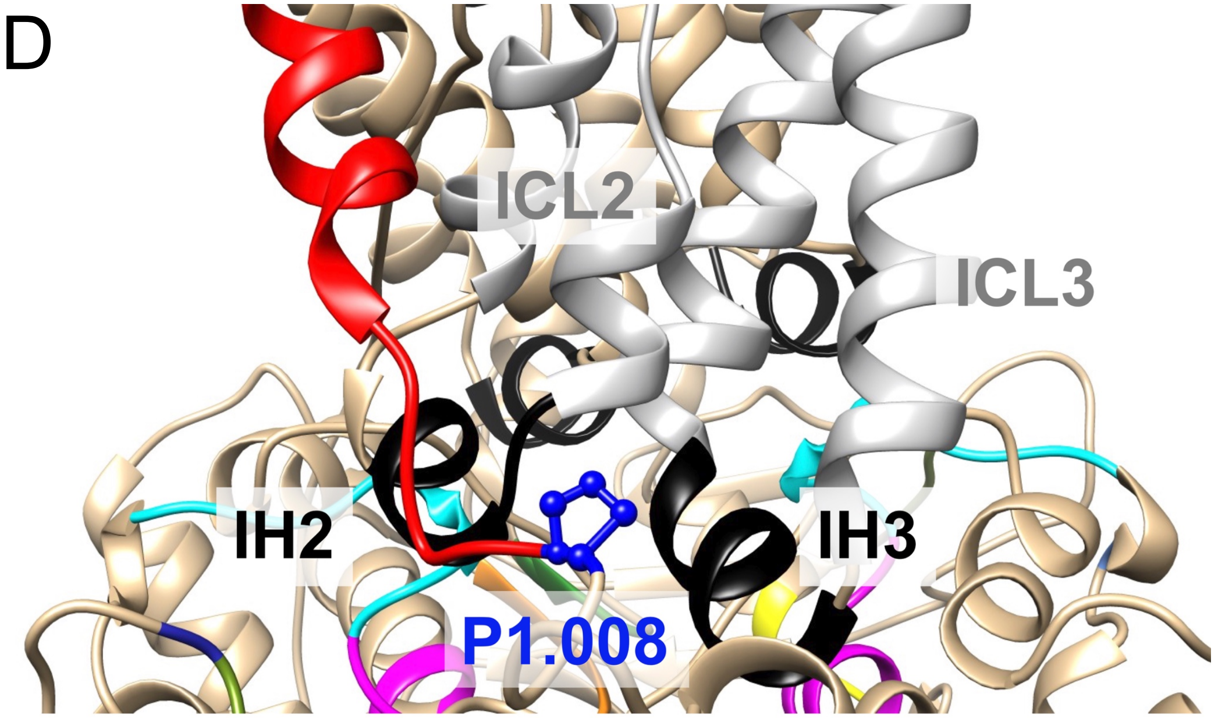


**Supporting Fig. 8: Conservation of the AtABCB1 P1.008 in CFTR/ABCC7 and speculative model on the role of P1.008 during ABCB1 regulation by TWD1.** Surface models of AtABCB1 (**A**) and HsCFTR/ABCC7 (**B**) revealing surface exposure of P1.008 and P1.184, respectively. (**C**) P1.008 is conserved between AtABCB1 and CFTR/ABCC7 and AtABCC1/2. (**D**) P1.008 lies in between intracellular/coupling helices 2 and 3, IH2 and IH3, respectively, establishing the connection between intracellular loops, ICL2 and ICL3, and the NBD2. Further, P1.008 is in direct connection with an alpha-helix (in red) leading to TMH12 shown to be involved in IAA binding (Bailly *et al.* 2012). Colour code of functional domains can be deduced from Fig. 1.
